# Droplet microfluidic sequencing of HIV genomes and integration sites

**DOI:** 10.1101/2020.09.25.314120

**Authors:** Chen Sun, Leqian Liu, Liliana Pérez, Xiangpeng Li, Yifan Liu, Peng Xu, Eli A. Boritz, James I. Mullins, Adam R. Abate

## Abstract

Sequencing individual HIV-proviruses and their adjacent cellular junctions can elucidate mechanisms of infected cell persistence *in vivo*. Here, we present a high throughput microfluidic method to sequence entire proviruses in their native integration site context. We used the method to analyze infected cells from people with HIV on suppressive antiretroviral therapy, demonstrating >90% capture and sequencing of paired proviral genomes and integration sites. This method should enable comprehensive genetic analysis of persistent HIV-infected cell reservoirs, providing important insights into the barriers to HIV cure.

---

Nearly 37 million people are infected with HIV worldwide, and despite huge research and clinical investments these infections remain incurable. A central obstacle to curing HIV lies within its mechanism of infection: HIV integrates its genome into the genome of infected cells. While many infected cells actively express virus genes, some infected CD4 T cells enter a state of reversibly nonproductive latent infection. Although antiretroviral therapy (ART) can suppress virus replication to undetectable levels and prevent progression to acquired immunodeficiency syndrome (AIDS), it lacks activity against latent cellular reservoirs^1-4^. Therefore, ART must be continued indefinitely to prevent virus rebound and recrudescence of disease progression. Lifelong ART is costly, has potential for toxicity, can be difficult to access, and fails to address HIV-associated stigma^5,6^. Thus, eliminating or suppressing HIV-infected cellular reservoirs to achieve a functional cure is an important goal of HIV research.

To elucidate how infected cellular reservoirs persist, paired analysis of integration sites and full-length provirus sequences is important. Identifying proviruses with matching integration sites has demonstrated that much of the latent reservoir consists of expanded cellular clones^7-12^. In ART-treated people living with HIV, lethally mutated proviruses far outnumber genetically intact ones capable of encoding replication-competent viruses^13,14^, the latter of which are difficult to identify and assess^15^. Sequencing the entire provirus is thus critical to determining whether it could give rise to rebound viremia after ART interruption. In addition, host genetic context may determine the expression level of a given provirus, thus influencing the stability of the host cell^10,15^. Importantly, understanding the relationships among provirus sequence integrity, clonality, and integration site effects requires that these multiple measurements be made for enough proviruses to faithfully represent complex cellular reservoirs *in vivo*.

Simultaneous assessment of genomes and integration sites has been hindered by the challenge of amplifying these components from individual DNA molecules. Current strategies distribute replicate aliquots of thousands of cell DNA-equivalents in microtiter plate wells at limiting dilution of the HIV proviruses, and then perform multiple displacement amplification (MDA)^15,16^. Wells containing proviruses are detected by subgenomic PCR, and additional MDA product is amplified again to recover near-full-length provirus genomes and products spanning virus-host integration sites, all of which are then sequenced. However, because the target provirus exists as a single copy within thousands of human genome equivalents, many reactions are required. In addition, due to a large background of human DNA during MDA, and the complexity of the PCRs to amplify the HIV genome and virus-host junctions, the reactions are prone to artifact generation, yielding products that contain spurious deletions, inversions, and HIV-human junctions that can confound analyses. Moreover, because multiple primer sets are required to amplify the genetically diverse HIV genome, the approach is biased towards variants that best match the primers and the regions targeted^14,17,18^. Most importantly, such methods are poorly suited to the even more extreme rarity of infected cells during ART, requiring massive expenses in time and resources to yield sequences for just a handful of proviruses. Consequently, while performing such analyses on many patients could elucidate important information about the cellular reservoirs of HIV, doing so with current technologies is impractical. To enable comprehensive HIV reservoir genomic characterization, a method that reliably and cost-effectively isolated and sequence rare proviruses *ex vivo* would be beneficial.

We describe a novel approach to characterize proviruses and their cellular genomic context within HIV reservoirs: Simultaneous Integration site and Provirus sequencing (SIP-seq). SIP-seq uses whole genome amplification in microfluidic droplets to amplify the HIV genome in its native context, and TaqMan PCR to tag droplets containing proviruses for sequencing. The result is the first technology providing the full-length virus genome connected to its host cell junctions in a single contiguous assembly. The speed and efficient reagent usage of droplet microfluidics allow recovery of single provirus genomes in a 150 million-fold higher background of DNA^19-21^. Using SIP-seq, we comprehensively profile the provirus population in multiple ART-treated people, to expand our understanding of the latent HIV reservoir. While we focus on HIV, our approach is applicable to viruses that insert into their host’s genome and thus provides a general platform for characterizing the genetics of diverse infections.

## Results

### Overview

Under effective ART, HIV replication can be suppressed to undetectable levels in the blood, but persist in rare infected cells in a reversibly latent form. Accurate quantitation of replication-competent proviruses is challenging, but recent estimates are as low as 1 in 100,000 or more CD4 T cells^8,22^. However, if therapy is stopped, viremia almost always rebounds to pre-therapy levels. CD4 T cells represent a critical reservoir for HIV^1^ and, thus, are the cells we interrogate for HIV persistence (Fig. 1a). The goal of our method is to recover from a population of millions of CD4 T cells all DNA fragments containing HIV genomes, and to individually sequence these molecules and their immediately adjacent host junctions (Fig. 2a). To isolate integrated HIV genomes, large DNA fragments are extracted from negatively selected or cultured CD4 T cells and encapsulated in microdroplets^20^. Each genomic fragment is then non-specifically amplified via MDA^23^, yielding enough product of each single provirus for sequencing (Fig. 1b, left). Next, multiplexed TaqMan PCR is used to identify and microfluidic sorting is used to isolate individual droplets containing HIV proviruses (Fig. 1b, right). Each positively sorted droplet is barcoded and sequenced. Lastly, reads from each droplet are mapped to a HIV reference genome and chimeras having host and virus sequences are detected to identify integration sites (Fig. 1c).

**Figure 1.**
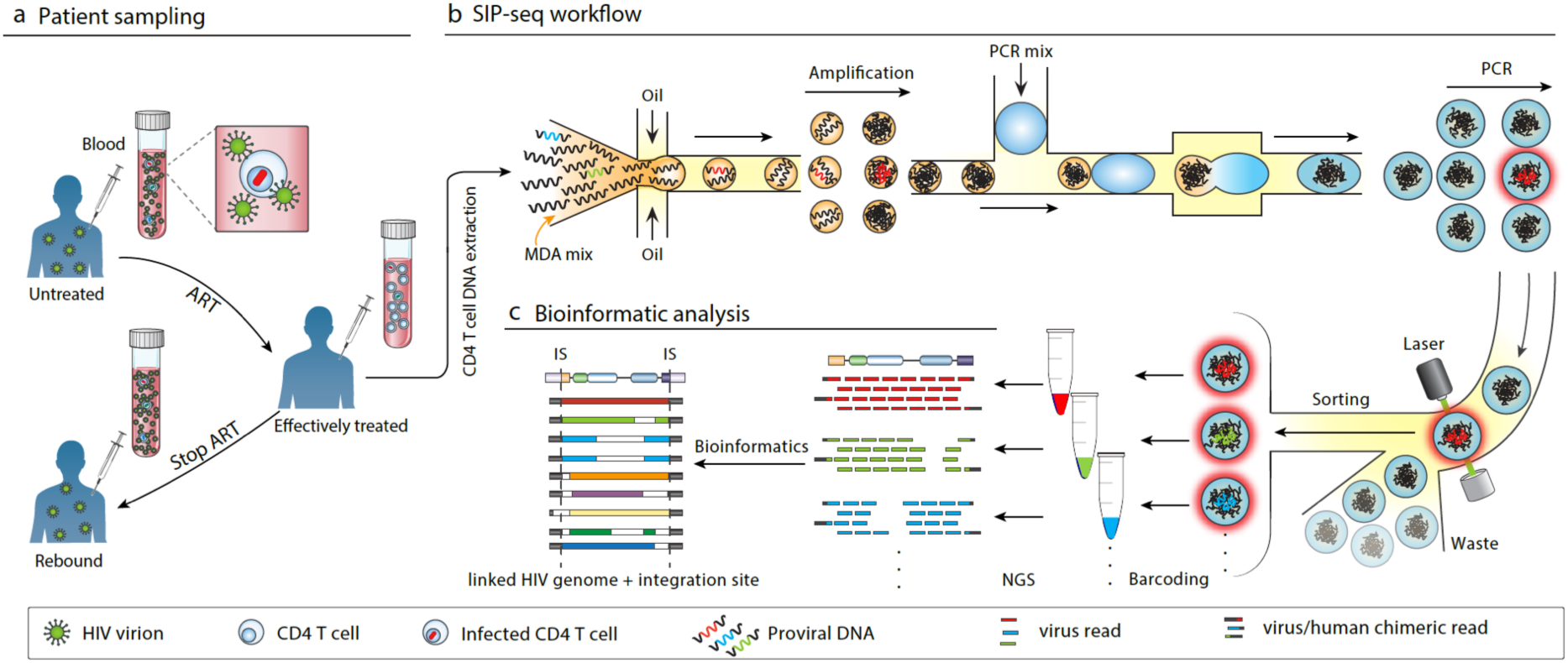
Application of SIP-seq to ART-treated individuals. (**a**) To study HIV persistence during ART, we sample CD4 T cells from effectively treated participants. (**b**) CD4 T cell DNA samples are processed with SIP-seq to generate single provirus sequencing libraries. gDNA is extracted and compartmentalized in droplets. DNA is MDA-amplified in each droplet, followed by PCR detection and sorting of HIV-positive droplets. DNA from each sorted droplet is barcoded and sequenced. (**c**) Sequence data is mapped to a reference HIV genome for simultaneous determination of individual virus genomes and integration sites.

**Figure 2.**
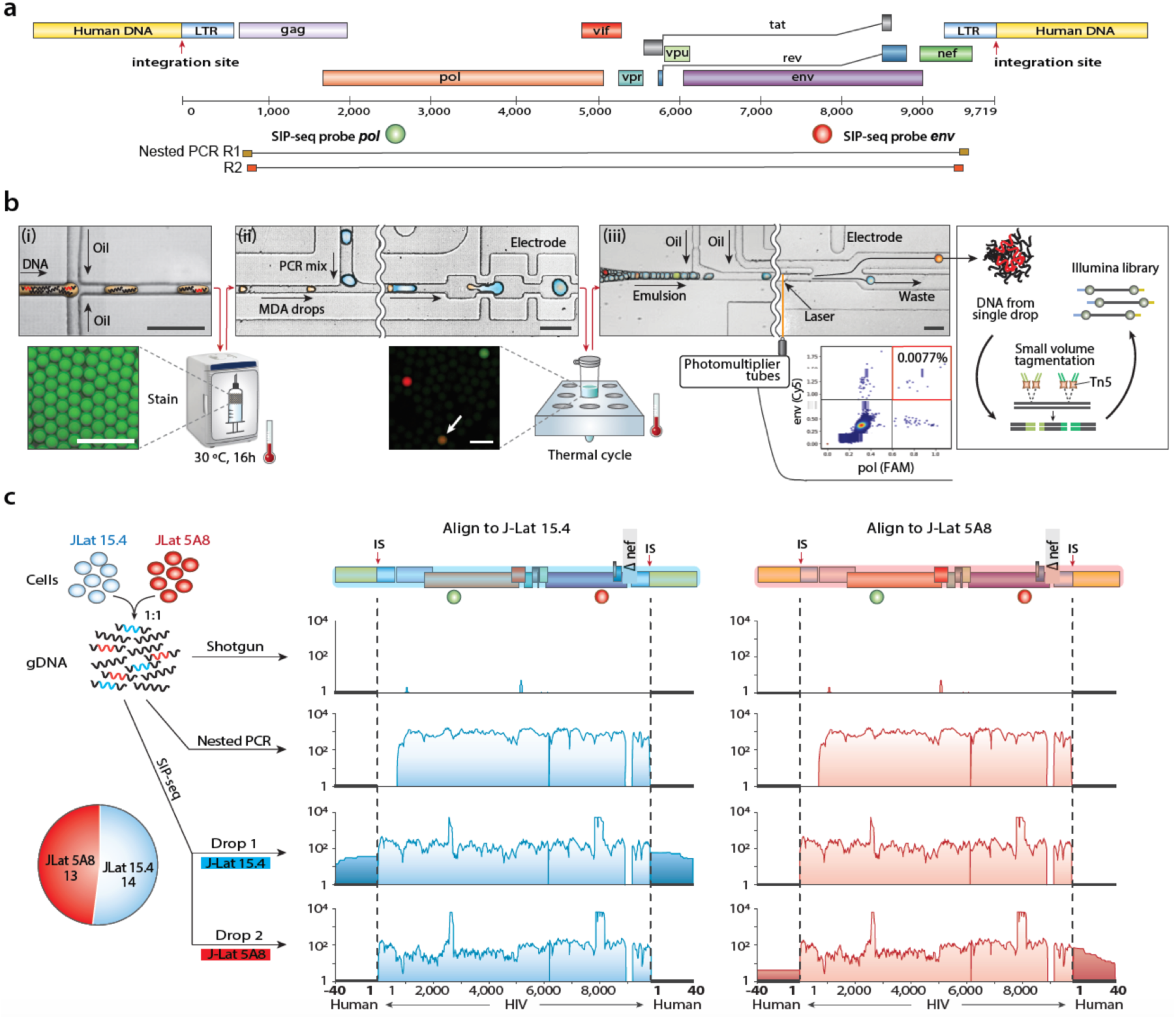
Demonstration and validation of SIP-seq with HIV-infected cell line samples. (**a**) TaqMan assays targeting the *pol* (green) and *env* (red) genes were used by SIP-seq to detect droplets containing HIV genomes. In comparison, nested PCR amplified near-full-length HIV genomes, but could not simultaneously recover the genome and flanking host sequences. Primer binding sites are shown relative to the schematic of an HIV genome integrated into a human genome. (**b**) SIP-seq microfluidics to identify HIV genome sequences, comprising: (i) encapsulation and amplification of DNA via MDA. An aliquot was stained with Eva Green for visualization; (ii) droplet merging to add TaqMan PCR reagents and in-droplet PCR. Fluorescence image shows representative drops after thermocycling indicating a double positive that would be isolated by sorting; and (iii) droplet sorting to select double PCR positives, with representative scatterplot of *pol* (FAM dye) and *env* (Cy5 dye) intensities. DNA from each sorted drop was processed for sequencing. Scale bars are 100 μm. (**c**) SIP-seq recapitulated HIV virus genomes and integration sites from a mixture of DNA from two J-Lat cell lines. gDNA from a mixture of the cell lines was processed with shotgun sequencing, sequencing of nested PCR products, and two SIP-seq drops containing either of two J-Lat proviral sequences. Both J-Lat proviruses had a deletion in the *nef* gene and sequences mapping to the full-length HXB2 genome; thus, no reads mapped to this region. The pie chart indicates recovery of genomes from a 50:50 mixture of DNA from the two cell lines.

Individual provirus encapsulation is necessary to obtain single provirus data. Our approach minimizes multiple virus encapsulation by partitioning cellular DNA to 1 provirus per >10,000 drops, yielding a doublet rate of below 0.01%. By contrast, brute force dilution approaches typically target 1 provirus per 3 wells, yielding a ∼17% doublet rate^15,16,24^. Thus, by leveraging the small volumes of droplet microfluidics, SIP-seq decreases reagent use by 100-fold, while also reducing doublets. Moreover, a unique and valuable property of performing the reactions in droplets is that they allow sequencing of picograms of DNA; this allows direct sequencing of the whole genome amplfication products without the need for additional complex multi-primer amplfication, yielding large, gapless contiguous assemblies that encapsulate the entire provirus genome connected to the integration site. This provides unambiguous information about virus genome completeness and integration site that is crucial to assessing the cellular reservoir.

### Provirus detection with SIP-seq

Random DNA cleavage and common deletions may result in partial genomes within a given droplet. To increase specificity for full-length proviruses, we employ a dually specific multiplexed TaqMan PCR targeting conserved regions of HIV *pol* and *env* spaced >5 kbp apart (Fig. 2a). Dual positive droplets in a representative HIV-infected cell sample were present at ∼1 in 13,000 and each recovered droplet yielded ∼3 pg of DNA. This amount of DNA was sufficient for sequencing. To confirm successful recovery of HIV proviruses, a small portion of the sorted DNA from each droplet was subjected to a TaqMan reaction targeting a different locus of the HIV genome (Supplementary Fig. 1).

### Microfluidics of SIP-seq

The microfluidics of SIP-seq consist of three devices: a droplet encapsulator, merger, and sorter. The encapsulator loads human genomic DNA fragments with MDA reagents in ∼20 μm droplets (Fig. 2b(i)). It runs at ∼20 kHz, allowing encapsulation of ∼10 billion DNA fragments ∼75 kbp in length in ∼20 million separate droplets over ∼15 min; each droplet thus contains ∼500 distinct fragments. The droplets are incubated at 30 °C to enable non-specific amplification before combining with TaqMan PCR reagents in the merger device. This device merges each MDA droplet with a ∼40 μm droplet containing TaqMan reagents, running at ∼2.5 kHz, and thus takes a few hours to process all droplets (Fig. 2b(ii)). Merged droplets are thermocycled for the TaqMan assay, yielding two-color fluorescence when an HIV genome is present (Fig. 2b(ii), lower). To isolate fragments containing HIV genomes, the droplets are introduced into a sorter operating at ∼1 kHz. If a droplet has fluorescence falling within specified gates, it is recovered via dielectrophoretic sorting and diverted into a collection tube (Fig. 2b(iii)). This is repeated for all detected HIV genomes, collecting each into a separate tube. All tubes are then prepared for sequencing^25^.

### Validation of SIP-seq on HIV cell lines

To validate SIP-seq, we used J-Lat 5A8 cells, a Jurkat-derived line containing full-length HIV with a frameshift in *env* and GFP in place of *nef*. As a comparator, we used conventional nested PCR to amplify the ∼9 kbp near full-length HIV genome. The SNPs detected by SIP-seq agreed with those detected by nested PCR with 0.02% mismatch likely due to MDA or PCR error (Supplementary Fig. 2). We also extracted virus-human chimeric reads from the SIP-seq data, and confirmed the known insertion site for this cell line^26^. This illustrates that SIP-seq can detect HIV SNPs and insertion sites.

To further assess the ability of SIP-seq to recover complete viral and flanking sequences, we performed a two-cell population experiment on DNA from J-Lat lines mixed at equal ratio (Fig. 2c, left), analyzed using conventional shotgun and PCR methods, and SIP-seq (Fig. 2c, middle and right). These cell lines (J-Lat 15.4 and 5A8) have identical virus genomes but different integration sites^26,27^. As expected, due to the small size of the HIV genome relative to that of the human genome, shotgun sequencing of all DNA detected no virus sequences (Fig. 2c, 1st row). Nested PCR resulted in excellent coverage of the near full-length virus genomes, but excluded a portion of the virus genome and integration site (Fig 2c, 2nd row). By contrast, SIP-seq provided a connected contig comprising the full-length proviruses and insertion sites (Fig. 2c, 3rd and 4th rows). The peaks at *pol* and *env* derived from abundant TaqMan PCR amplicons used for droplet detection. Reads from two representative drops mapped to either of the two cell lines with minimal cross contamination. In total, we sequenced 27 DNA fragments containing single virus genomes and integration sites. Moreover, the detection rates of the cell lines matched the known input ratio, demonstrating no bias in recovery.

### Analysis of HIV in CD4 T cells from an ART-treated person after *in vitro* expansion

Infected cell expansion (ICE) from *ex vivo* samples allows rare infected CD4 T cells to be diluted and cultured, such that 1/10 to 1/3000 cells corresponds to an infected clone post expansion^28^. This reduces infected cell rarity and enables downstream studies. Moreover, ICE cultures provide an excellent test of SIP-seq, since they contain proviruses from HIV-positive people in their original host genome contexts. The cells are prepared by plating resting CD4 T cells from an ART treated person at 1 infected cell per ∼5 wells (100-3,000 total cells per well), followed by stimulation and *in vitro* culture to allow proliferation (Fig. 3a). We extracted and analyzed DNA from several wells of an ICE culture plate using all three methods (shotgun, nested PCR, and SIP-seq). As expected, due to the rarity of infected cells even with ICE enrichment, shotgun sequencing recovered little of the HIV genomes (Fig. 3b, top row). By contrast, both SIP-seq and nested PCR yielded excellent coverage of the virus genomes (Fig. 3b, rows 2 and 3), with SIP-seq providing the insertion sites in host gene ARIH2 (Fig. 3b, row 3). Notably, a second well from an ICE culture was also examined, in which a single HIV LTR region PCR was positive (Supplementary Fig. 3). In this case, neither shotgun nor nested PCR yielded coverage of the HIV genome (Fig. 3c, rows 1 and 2). SIP-seq, by contrast, yielded an HIV genome harboring an inversion and large deletion with insertion in host gene RABL6 (Fig. 3c, row 3). We confirmed this result using custom primers targeting the adjacent human sequences (Figs. 3b and 3c, bottom panels). Thus, SIP-seq is well suited to characterizing the full genetic diversity of infected cells *in vivo*.

**Figure 3.**
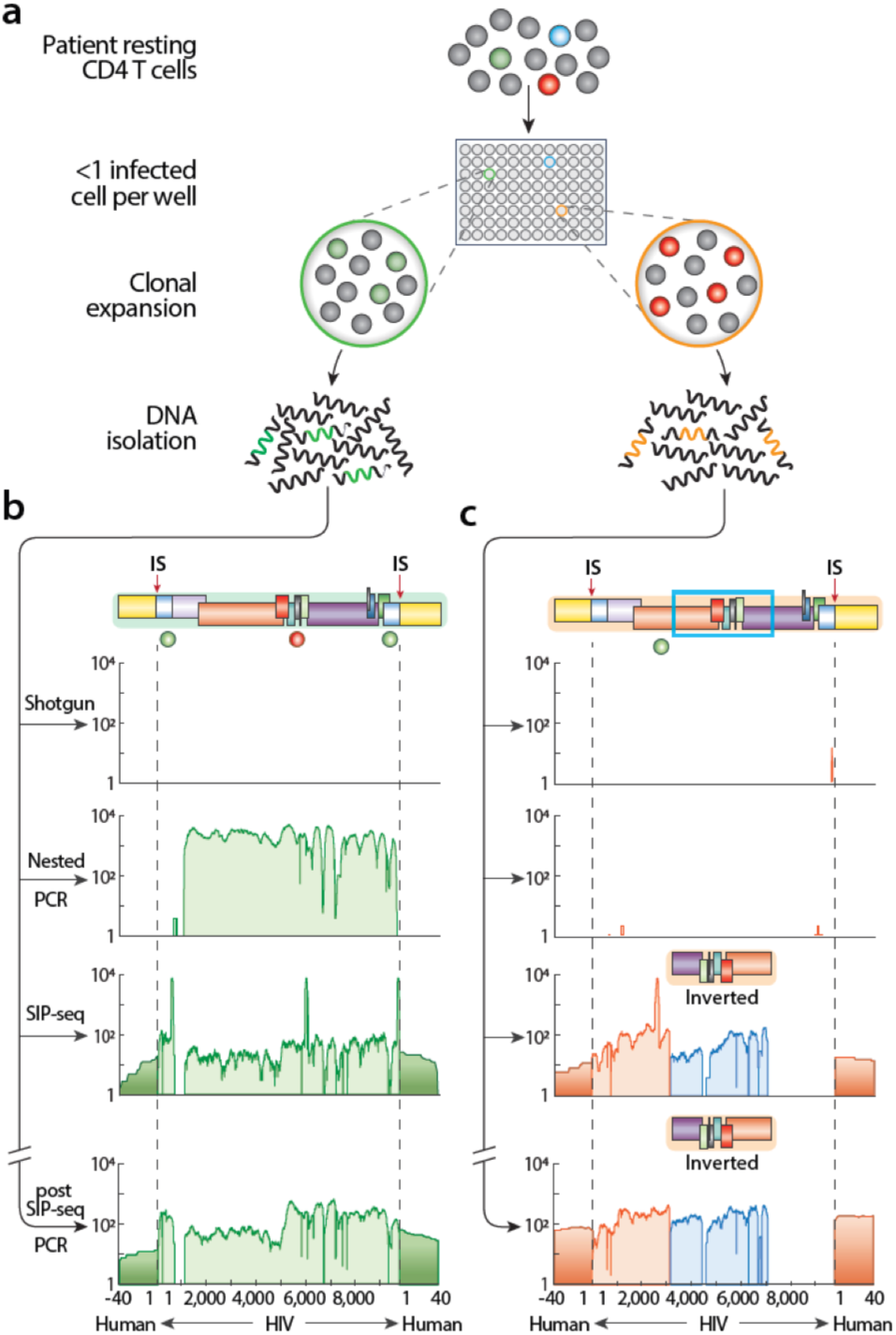
SIP-seq of HIV in ART treated participant CD4 T cells after clonal expansion. (**a**) Infected cell expansion (ICE) was prepared by seeding participant CD4 T cells to < 1 infected cell per 5 wells and culturing for clonal expansion. Two ICE clones were separately processed to sequencing with different technologies. (**b**) SIP-seq detected an intact HIV genome integrated into a human genome. (**c**) SIP-seq identified an integrated HIV genome containing an inverted sequence and large deletion. Nested PCRs using primers designed based on the SIP-seq results confirmed the HIV genomes (bottom panels of **b** and **c**).

### Genomic landscape of HIV infection in ART treated individuals

We next characterized the genomic landscape of HIV proviruses isolated directly from the CD4 cells of infected persons (Fig. 4, Supplementary Table 1). Individuals #1 and #2 were receiving effective ART with virus RNA loads in blood plasma below 20 copies/mL at the time of study. In these two individuals, we recovered 29 provirus genomes and integration sites (Fig. 4a, b). None of these proviruses were intact. Of the recovered genomes, 14 yielded integration sites, with 4 having junctions on the 3’ and 5’ ends. Two had intact *gag* and *pol* and were integrated into host genes STAT5B and HIVEP1 of participant #1 in the opposite orientation relative to the gene. We also observed higher insertion in genic over intergenic (12 versus 2) regions, with no preference in orientation relative to host genes (7 same vs 7 opposite) (Fig. 4c(i)). Sixteen proviruses had large deletions that rule-out replication-competency, though not necessarily excluding virus protein expression and resulting cellular activation and immune recognition^8,29^. Proliferation of latently infected cells is thought to play an important role in HIV persistence^10,12,30^. Our analysis reveals three clonal lineages that likely originate from separate infected cells (Fig.4c(ii)) and several proviruses that matched sequences obtained from previous studies of the same individuals^31^ (Supplementary Fig. 4). Our results thus support the clonal origin of these lineages and provide novel integration site information.

**Figure 4.**
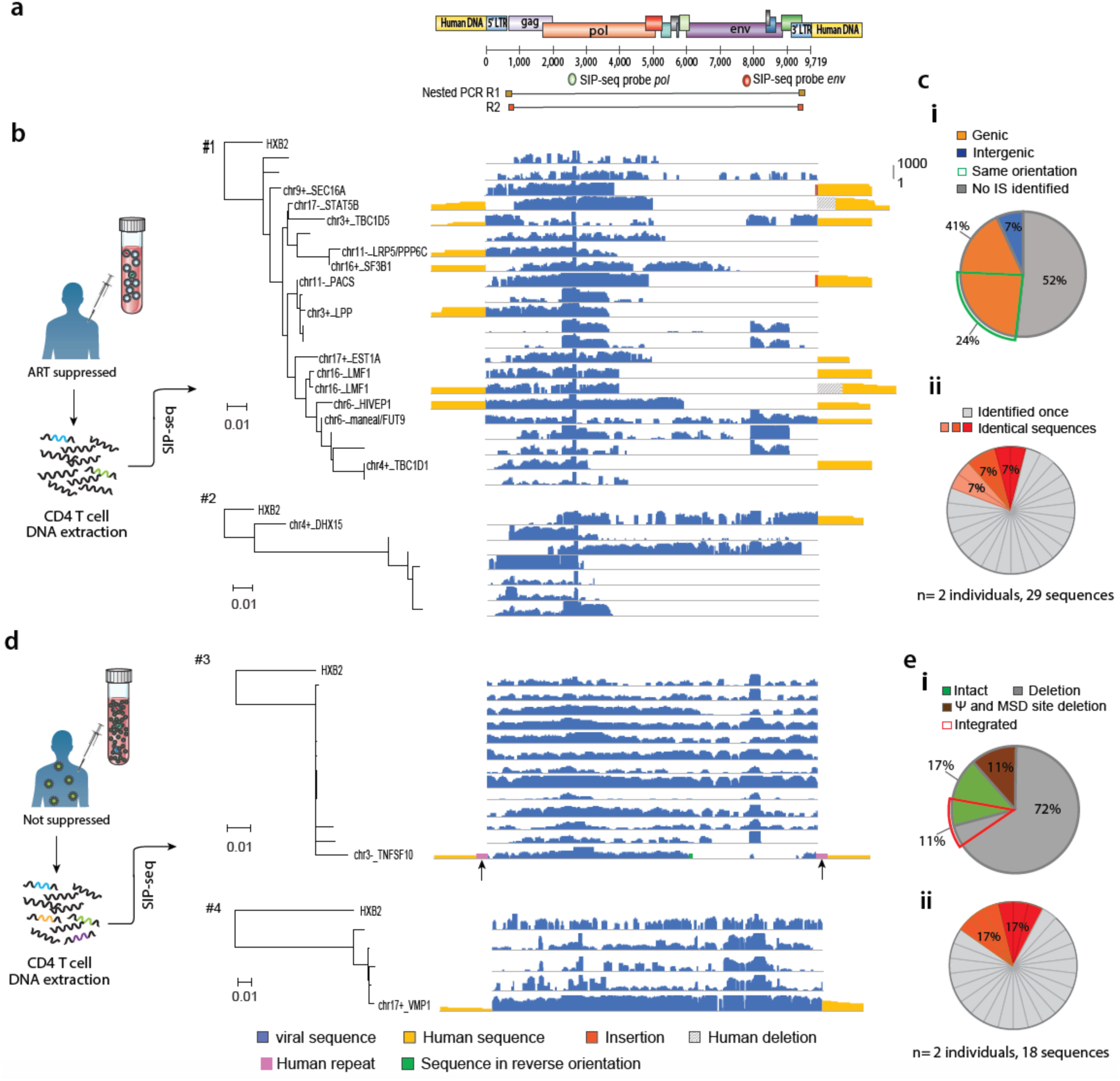
SIP-seq of HIV proviruses and flanking sequences from CD4 cells directly from infected individuals. (**a**) Schematic showing positions of TaqMan PCR probes in HIV *pol* and *env*, and locations of nested PCR primers used for near-full-length genome amplification. (**b**) SIP-seq of DNA from CD4 T cells from individuals (#1 and #2) with ART suppression. Phylogenetic trees were generated using a ∼500 bp region of *pol* from all HIV sequences obtained from each participant. Host genes and provirus orientation relative to the host gene for each sequence are indicated unless no integration site was identified. Each bar represents a coverage map of an individual HIV genome and its integration site. (**c**) Integration sites were not random (**i**) and expanded clones were found (**ii**) in the participants on ART suppression. (**d**) SIP-seq of DNA from CD4 T cells from donors without ART suppression (#3 just started ART and #4 had ART suspended) (**e**) Intact and defective HIV (**i**) and expanded clones (**ii**) in participants #3 and #4.

Importantly, >90% of the obtained proviruses contained deletions in the primer positions used for most near-full-length nested PCRs, and would thus not have been characterized effectively by those methods^14,17,32^. Furthermore, we recovered genome variants with mutations in LTR regions not captured by nested PCR. For Participant #1, 4 of 22 sequences contained intact 5’LTR through the major splice donor and thus have the potential to drive transcription of LTR transcripts into cellular sequences (Supplementary Fig. 5).

At the time of sampling, individual #3 had just started ART and # 4 received ART for a year before going off therapy for five years and both had viral loads >100,000 copies/mL (Fig. 4d). As expected for the setting of active HIV replication, SIP-seq revealed abundant unintegrated and intact virus compared to the two individuals receiving effective ART^33,34^. Of the 18 sequences characterized, 16 were unintegrated, two of which were intact (Fig. 4e(i)). Only one provirus from each individual (one intact, one defective) was integrated, yielding integration sites. In Participant 3, one provirus had a large internal deletion flanked by a direct repeat of ∼124 bp of host sequences derived from the gene TNFSF10 (indicated by arrows). In Participant 4, an intact provirus was integrated into the VMP1 gene. We also found two groups of three sequences likely originating from clonally proliferating cells (Fig.4e(ii)). Proviruses from Participant 3 had duplications of the transcription factor TCF-1α binding site in several LTRs (Supplementary Fig. 6).

## Discussion

SIP-seq allows study of HIV proviruses that could not have been recovered by current PCR methods alone, and provides a more comprehensive analysis of the HIV genetic landscape *in vivo*. Capturing full-length proviral HIV genomes with their associated integration sites is essential for characterizing the latent reservoir and its contribution to HIV persistence. However, the rarity and lack of distinct surface markers for latently infected cells pose major obstacles to current techniques. While microfluidic enrichment and pooled sequencing methods allow characterization of rare genome sequences^20,36^, SIP-seq is the first technology to provide robust sequencing of individual rare proviruses in a format that is fast, cost effective, and scalable.

A limitation of SIP-seq is that it requires short-ranged TaqMan PCR amplification to detect DNA fragments with integrated provirus. If the targeted regions are not present in a provirus, it will not be detected. For this reason, we designed primers to target conserved regions of HIV genomes, ensuring that proviruses with the potential for infectivity will be enriched for detection. Moreover, SIP-seq is flexible with respect to the TaqMan primers used, allowing their optimization to capture unexpected variants. We can also use established intact proviral DNA assay (IPDA) primers in SIP-seq to selectively sequence intact proviruses^13^. Additionally, this minimum requirement is less constraining than nested PCR, which requires multiple primer pairs to amplify an incomplete portion of the HIV genome^17,32^. As we have shown in a case example, viruses in which these regions are absent or divergent can easily be missed. Moreover, the multiple rounds of PCR used to separately amplify the provirus genome and its integration site after microwell MDA have the potential to generate artifacts^37,38^. Because SIP-seq yields contigs containing the full-length provirus physically connected to its integration site on both sides, the resultant sequences are of high confidence.

By applying SIP-seq to people receiving ART, we confirm that proviruses in people with undetectable viremia are most often defective and found in clonally expanded cells, and that ongoing viremia is associated with higher levels of unintegrated and intact virus genomes. We also made several novel observations. Most importantly, >90% of the proviruses we observed would not have been recovered by nested PCR methods commonly in use. These proviruses were defective but can still be immunogenic and influence cell function^8,29^. These findings demonstrate that SIP-seq is a valuable tool for rigorous assessment of the latent reservoir, to confirm hypothesized attributes and discover novel ones. While we focused on HIV-1 infection, SIP-seq should be applicable to other retrovirus infections that include an integrated state as part of their life cycle.

## Methods

### Microfluidic device fabrication

The microfluidic devices were fabricated by soft lithography. Photomasks designed in AutoCAD were printed on transparencies. The features on the photomask were transferred to a negative photoresist (MicroChem, SU-8 2025) on a silicon wafer (University Wafer) by UV photolithography. Polydimethylsiloxane (PDMS, Dow Corning, Sylgard 184) prepolymer mixture was poured over the patterned silicon wafer and cured in a 65 °C oven for 2 h. The PDMS replica was peeled off and punched for inlets and outlets by a 0.75 mm biopsy core (World Precision Instruments). The PDMS slice was bound to clean glass by oxygen plasma cleaner (Harrick Plasma), followed by baking at 65 °C for 1h to ensure strong bonding. The microfluidic channels were treated with Aquapel (PPG Industries) and baked at 65 °C for 30 min for hydrophobicity.

### JLat cell culture and genomic DNA extraction

JLat HIV latency clones 5A8 and 15.4 were kindly provided by Dr. Mauricio Montano at the Gladstone Institute at UCSF^39^. JLat 5A8 and JLat 15.4 cells were grown in RPMI 1640 media (Gibco #11875093) supplemented with 10% fetal bovine serum (FBS, Gibco #26140079), and penicillin-streptomycin (Gibico #15140122). Cells were incubated with 5% CO_2_ at 37°C. The genomic DNA of JLat cells and CD4 T cells from individuals #3 and #4 were extracted with a Quick-DNA Miniprep Plus kit (Zymo Research #D4068) according to the product protocol. The cell culture and DNA extraction from ICE cultures (1737.H3 and TC1288.B9) were conducted in the Gladstone biosafety level 3 (BSL3) facility.

### Isolation of CD4 T cells

Participant recruitment and informed consent were performed under Institutional Review Board (IRB)-approved protocols at the US National Institutes of Health (NIH). Peripheral blood mononuclear cells (PBMC) were isolated from whole blood by density gradient centrifugation. PBMC were incubated with Fcγ-receptor-blocking reagent for 10 minutes and stained with LIVE/DEAD Aqua stain, CD3-APC-H7 (BD Biosciences; Cat# 641406), CD4-BV785 (BioLegend; Cat# 317442), CD8-PacBlue (Invitrogen; Cat# MHCD0828), CD14-BV650 (BioLegend; Cat# 301836), CD16-PerCP/Cy5.5 (BioLegend; Cat# 302028), CD19-BV605 (BD Biosciences; Cat# 562653), CD20-BV570 (BioLegend; Cat# 302332), CD27-Alx700 (BioLegend; Cat# 302814), CD32-PE (BioLegend; Cat# 303206); CD45RO-ECD (Beckman Coulter; Cat# IM2712U), CD123-PE/Cy5 (BD Biosciences; Cat# 551065), and TCRγδ-APC (BD Biosciences; Cat# 555718). CD4 T cells were isolated by fluorescence-activated cell sorting (FACS) on a FACSAria (Becton Dickinson) using previously described protocols^31^.

### Extraction of cell-associated DNA

Sorted cells were sedimented by centrifugation at 400 x *g* for 7 minutes at 4°C, lysed in RNAzol RT at <5 ⨯ 10^6^ cells/mL, homogenized by pipetting, and stored at −80°C until extraction. For DNA extraction, 0.4 volume of RNAzol RT of sterile H_2_O was added to each lysate to allow aqueous and organic phase separation. The organic phase of each lysate was solubilized in DNAzol (Molecular Research Centers) to allow DNA extraction according to the manufacturer’s instructions.

### Nested PCR

The nested PCR was performed as described previously^17^. Post SIP-seq nested PCR amplified the full-length HIV genomes and integration sites in two ICE clones. The primers used in post SIP-seq nested PCR were designed based on the adjacent human sequences and are listed in supplementary table 3. Amplified DNA was prepared for NGS using the Illunima Nextera XT DNA Library Preparation Kit according to the manufacturer’s instructions.

### MDA in microfluidic emulsion droplets

The MDA reaction mixture was prepared with phi29 DNA Polymerase (New England BioLabs catalog no. M0269L) following the manufacturer’s protocol. Purified T cell DNA was added into a 100 μl reaction containing 1X phi29 DNA Polymerase Reaction Buffer, 200 μg/ml BSA, 200 μM dNTPs, 25 μM random hexamers and 60 U phi29 polymerase. The reaction mixture was loaded immediately into a 1 ml syringe backfilled with HFE-7500 fluorinated oil (3M, catalog no. 98-0212-2928-5) and injected into a flow-focus droplet maker (Supplementary Fig.7a) at a constant rate of 400 μl/h by a syringe pump (New Era). HFE-7500 oil with 2% (w/w) PEG-PFPE amphiphilic block copolymer surfactant (RAN biotechnologies, catalog no. 008-FluoroSurfactant-1G) was loaded into another 1 ml syringe and injected into the droplet maker at 1000 μl/h. The preparation of the MDA reaction mix and generation of droplets were done within 15 min to minimize amplification in bulk. Monodispersed droplets were generated and collected into a 1 ml syringe and incubated at 30 °C for 16 h. An aliquot of the MDA reaction was mixed with a DNA dye (EvaGreen, Biotium) before being emulsified as described above. After incubation, the syringe with MDA droplets was placed in a 65 °C oven for 3 min to deactivate phi29 DNA polymerase. The EvaGreen stained drops were examined by fluorescence microscopy (EVOS) to determine MDA efficiency.

### Droplet merging and digital droplet TaqMan PCR

MDA droplets and 1.125X TaqMan PCR mix were each injected into the microfluidic merger device as shown in Supplementary Fig.7b. 1X TaqMan PCR mix contained 1X Platinum Multiplex PCR Master Mix (Life Technologies, catalog no. 4464269), 200 nM TaqMan probes (IDT), 1 μM primers (IDT), 2.5% (w/w) Tween® 20 (Fisher Scientific), 2.5% (w/w) Poly(ethylene glycol) 6000 (Sigma-Aldrich) and 0.8 M 1,2-propanediol (Sigma-Aldrich). Tween® 20 and Poly(ethylene glycol) 6000 were used for droplet stability during thermal cycling. 1,2-propanediol was used as PCR enhancer for low temperature denaturation. TaqMan sets were used: *pol*-probe (HXB2 position 2586–2604), /56-FAM/AA GCC AGG A/ZEN/A TGG ATG GCC /3IABkFQ/; *pol*-F, GCA CTT TAA ATT TTC CCA TTA GYC CTA; *pol*-R, CAA ATT TCT ACT AAT GCT TTT ATT TTT TC; *env*-probe (HXB2 position 7791–7815), /5Cy5/CC ATA GTG C/TAO/T TCC TGC TGC TCC CAA /3IAbRQSp/; *env*-F, GGC AAR GAG AAG AGT GGT GCA; *env*-R, GYC TGG CYT GTA CCG TCA GC. MDA droplets were reinjected into the merger device at 50 μl/h. A stream of HFE-7500 oil with 2% (w/w) PEG-PFPE surfactant at a flow rate of 500 μl/h was used as spacer. PCR reagent (flow rate 200 μl/h) drops were formed in HFE-7500 oil with 2% (w/w) PEG-PFPE surfactant (flowed at 600 μl/h) and subsequently merged with the MDA drops pairwise. The merging of the two drops was achieved at the liquid salt electrode, which was connected to a cold cathode fluorescent inverter and DC power supply (Mastech) to generate a ∼2 kV AC potential at the electrode from a 2 V voltage. The merged drops containing MDA products and now 1X TaqMan PCR mix were collected into PCR tubes. The bottom oil layer was removed and replaced with FC-40 fluorinated oil (Sigma-Aldrich, catalog no. 51142-49-5) with 5% (w/w) PEG-PFPE surfactant to maintain emulsion stability during thermal cycling. The cycling was performed on a T100 thermal cycler (Bio-Rad) with conditions: 2 min at 86 °C; 35 cycles of 30 s at 86 °C, 90 s at 60 °C and 30 s at 72 °C; and finally, 5 min at 72 °C. Low denaturation temperature of 86 °C was used to minimize DNA fragmentation.

### Dielectrophoretic sorting

After thermal cycling, the drops were transferred to a 1 ml syringe and reinjected into a microfluidic DEP sorter (as shown in Supplementary Fig.7c) at 80 μl/h. HFE-7500 oil was loaded into a 5 ml syringe and injected into the DEP device as spacer oil at a flow rate of 1220 μl/h. Another stream of HFE-7500 oil at 1300 μl/h was introduced at the sorting junction to drive the drops into waste collection when the electrode is deactivated. A negative pressure from a syringe at −1300 μl/h was applied to the waste collection to make sure drops flow into waste when deactivated. The salt electrodes and moat were filled with 2M NaCl solution. The PMTs (Thorlabs, PMM01 model) were controlled by a data acquisition card and a LabVIEW program (National Instruments) to measure droplet fluorescence when passing the laser. The sorting electrode is activated when the fluorescence intensity is higher than a pre-set threshold. A high-voltage amplifier (Trek) was used to amplify the electrode pulse to 1 kV for DEP sorting. The sorted drops were collected into individual PCR tubes with one droplet per PCR tube for single viral sequencing.

### Library preparation and sequencing of single droplets

The sorted individual droplet with its carrier oil in a PCR tube was dried out in a vacuum chamber for ∼ 1.5 h. 1.5 μl DI H_2_O was then added to dissolve sorted DNA, of which 0.5 μl DNA was used to confirm sorting of the target by qPCR on a different region of the virus genome than *pol* and *env*. The qPCR was performed using 1X Kapa probe fast qPCR 2X master mix (KAPA Biosystems Inc, catalog no. KK4702), 140 nM TaqMan probe (IDT), 300 nM of each primers (IDT) and 0.5 μl sorted DNA with condition: 3 min at 95 °C; 40 cycles of 5 s at 95 °C, 15 s at 58 °C, and 30s at 60 °C in QuantStudio 5 Real-Time PCR System (Thermo Fisher Scientific). The sequences of the probes and primers are listed in supplementary table 2. The shifts in the curves (for one or more of the 5 loci) confirms sorting of HIV sequence-containing drops.

To construct Illumina libraries from picogram amounts of DNA recovered from sorted single drops, we slightly modified the standard Illunima protocol to increase reaction efficiency by reducing reaction volume and optimizing enzyme concentrations. The remaining 1μl re-dissolved contents were tagmented using 0.6 μl TD Tagmentation buffer and 0.3 μl ATM Tagmentation enzyme from Nextera XT DNA Library Prep Kit (Illumina, catalog no. FC-121-1030) for 5min at 55 °C. 1 μl NT buffer were added to neutralize the tagmentation. The tagmented DNA was then mixed with PCR solution containing 1.5 μl NPM PCR master mix, 0.5 μl of each index primers i5 and i7 from Nextera Index Kit (Illumina, catalog no. FC-121-1011) and 1.5 μl H_2_O, and placed on a thermal cycler with the following program: 3 min at 72 °C; 30 s at 95 °C; 20 cycles of 10 s at 95 °C, 30 s at 55 °C, and 30 s at 72 °C; and finally 5 min at 72 °C. The DNA library was purified and size-selected for 200-600 bp fragments using Agencourt AMPure XP beads (Beckman Coulter), and quantified using Qubit dsDNA HS Assay Kit (Thermo Fisher Scientific) and High Sensitivity DNA Bioanalyzer chip (Agilent). The library was sequenced using Illumina Miseq PE150 or Hiseq PE300.

### Bioinformatic analysis

Sequencing reads passing quality control were mapped to the HIV reference (HXB2) using Bowtie 2^40^. Genomic coverage as a function of genome position was generated using SAMtools^41^. ∼1 million paired-end reads of 150 bp were used for enriched or unenriched cell line samples, ∼2 million paired-end reads of 150 bp each were analyzed for patient derived ICE samples, and ∼10 million reads of 300 bp were used to study patient samples. BCFtools, within SAMtools, was used for variant calling. Chimeric reads and their soft-clipped regions from alignments (non-HIV regions) were extracted using extractSoftclipped (https://github.com/dpryan79/SE-MEI). Soft-clipped reads were analyzed by a web base tool for integration site localization (https://indra.mullins.microbiol.washington.edu/integrationsites/). Reads were *de novo* assembled using SPAdes^42^ and contigs evaluated by QUAST^43^ to confirm integration sites and provide additional information on proviral structures. Assembled sequences were grouped by participant and used to construct maximum likelihood trees in DIVEIN^44^ (https://indra.mullins.microbiol.washington.edu/DIVEIN/). HIV Highlighter plots of LTR regions for each participants were created using Highlighter^45^ at https://www.hiv.lanl.gov/content/sequence/HIGHLIGHT/highlighter_top.html.

## Acknowledgements

This work was supported by a Chan Zuckerberg Biohub grant to A.R.A., a National Institutes of Health grant (R01 HG008978) to A.R.A., a National Institutes of Health grant (U01 AI129206) to A.R.A. and E.A.B., and National Institutes of Health Grants (R01 AI125026 and R33 AI122361) to J.I.M. The content is solely the responsibility of the authors and does not necessarily represent the official views of the National Institutes of Health.

## Author Contributions

C.S. and A.R.A conceived of the project. C.S., L.L., X.L., Y.L. and P.X. performed the experiments. C.S. sequenced the samples and analyzed the data. C.S. and A.R.A. wrote the initial draft of the manuscript. L.P. assisted with patient sample processing. J.I.M. and E.A.B. revised the manuscript. All authors read, reviewed, and approved the manuscript.

