## Supplementary data for "Droplet microfluidic sequencing of HIV genomes and integration sites"

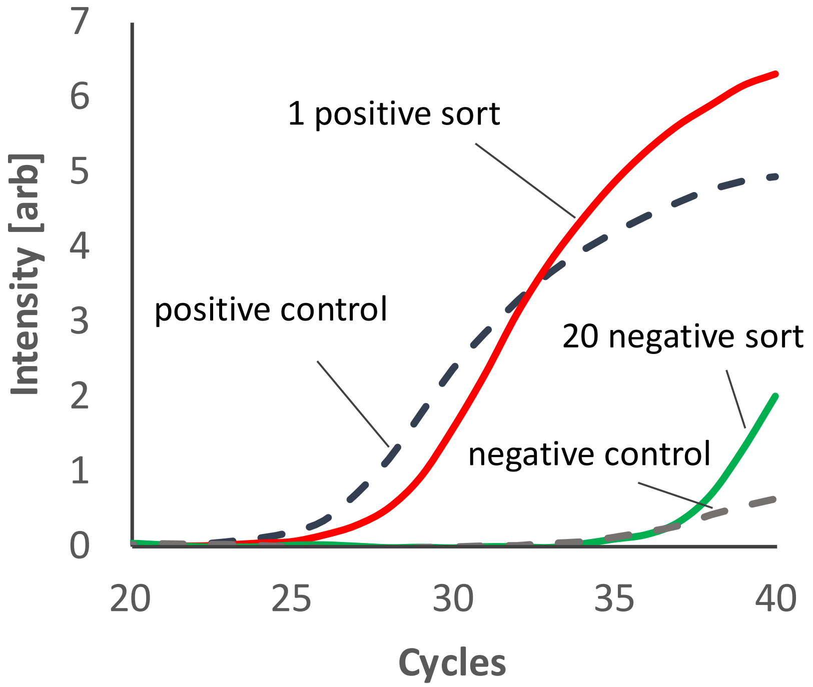


**Supplementary Figure 1.** qPCR detection of HIV sequence in sorted drops. DNA recovered from 1 positive drop or from 20 negative drops was subjected to a HIV-specific TaqMan PCR reaction. The qPCR curve for 1 sorted positive drop (red line) shifts to lower cycles compared to that for 20 negative drops (green line), illustrating successful recovery of HIV proviruses. The positive control used was JLat DNA and the negative control was uninfected Jurkat cell DNA.

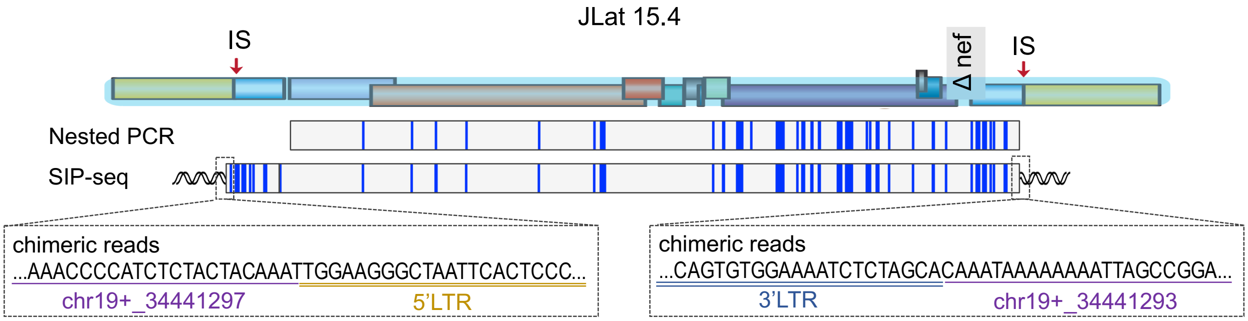


**Supplementary Figure 2.** SIP-seq detection of HIV SNPs and integration site of J-Lat 15.4 cell line. The SNPs detected by SIP-seq > 99.98% matches to nested PCR result. The integration site is determined by chimeric reads analysis and matches to the known integration site of this cell line[1]. The 4 nucleotide offset in 5' and 3' integration sites is consistent with the known process of duplication of cellular sequences during HIV integration. Vertical blue lines indicate mutations relative to an HIV-HXB2 reference sequence.

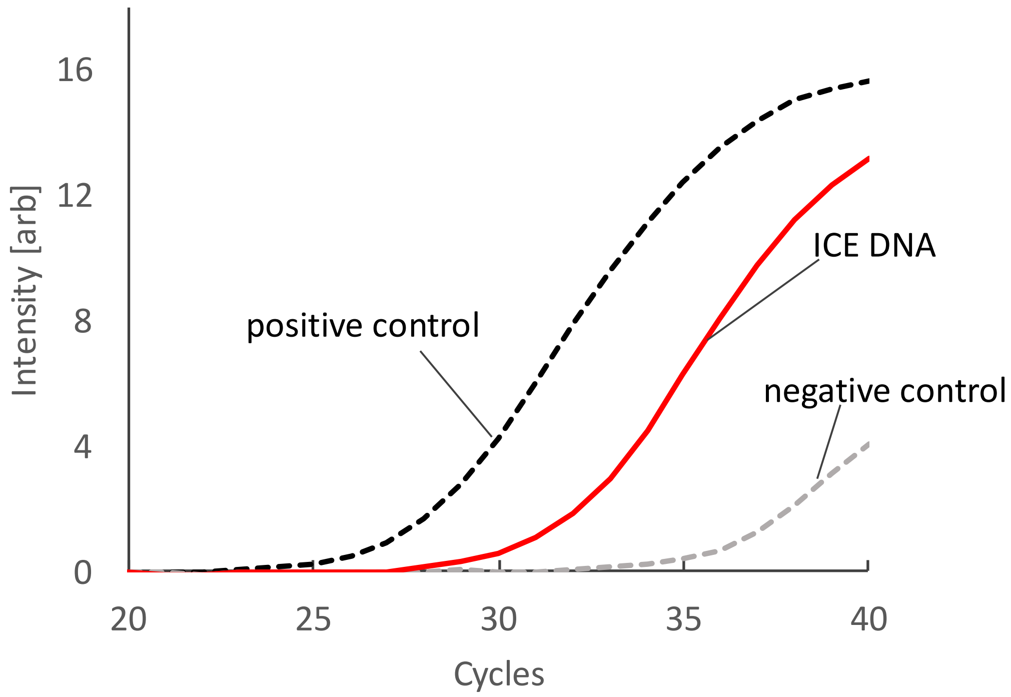


**Supplementary Figure 3.** qPCR detection of HIV sequence in an ICE culture. The ICE culture well contained HIV sequence confirmed by a HIV LTR region PCR. The positive was JLat DNA and negative control was Jurkat DNA.

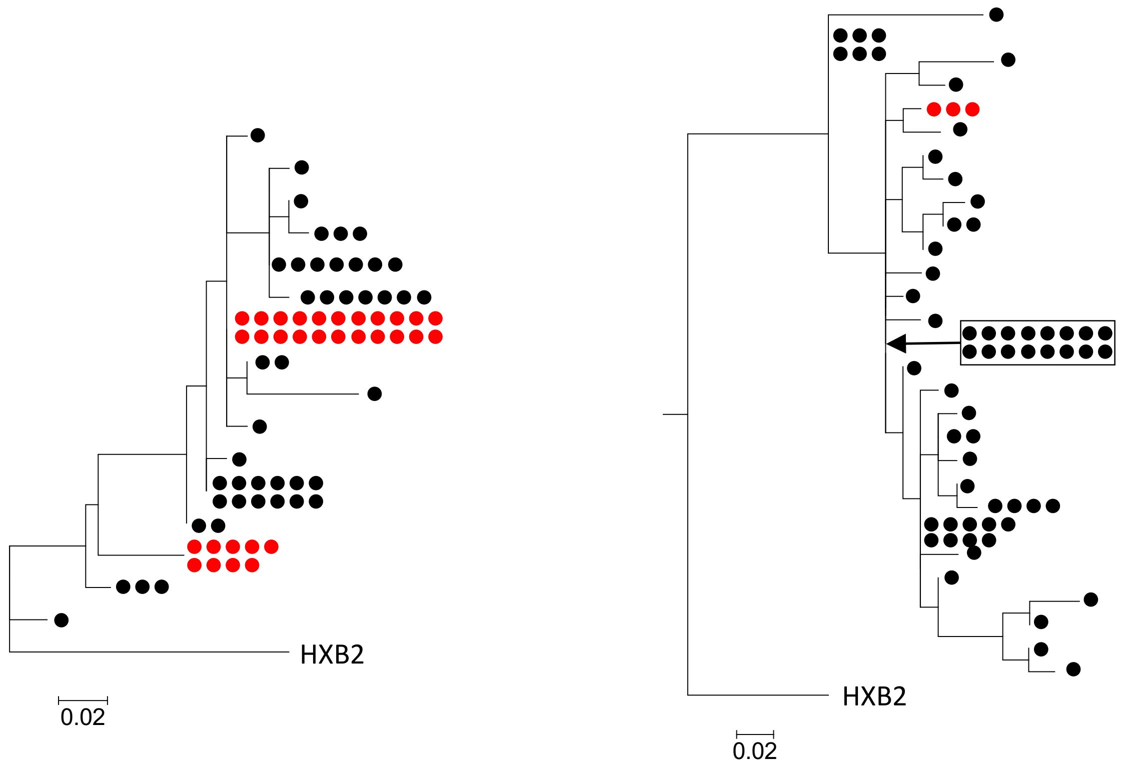


**Supplementary Figure 4.** Mapping of sequences detected by SIP-seq to previous phylogenetic trees made by PCR amplification of *env* from the same donor samples (Left: individual #1, right: individual #2). Red dots are sequences in the trees that one of SIP-seq sequences matched and black dots are sequences that no SIP-seq sequences matched.

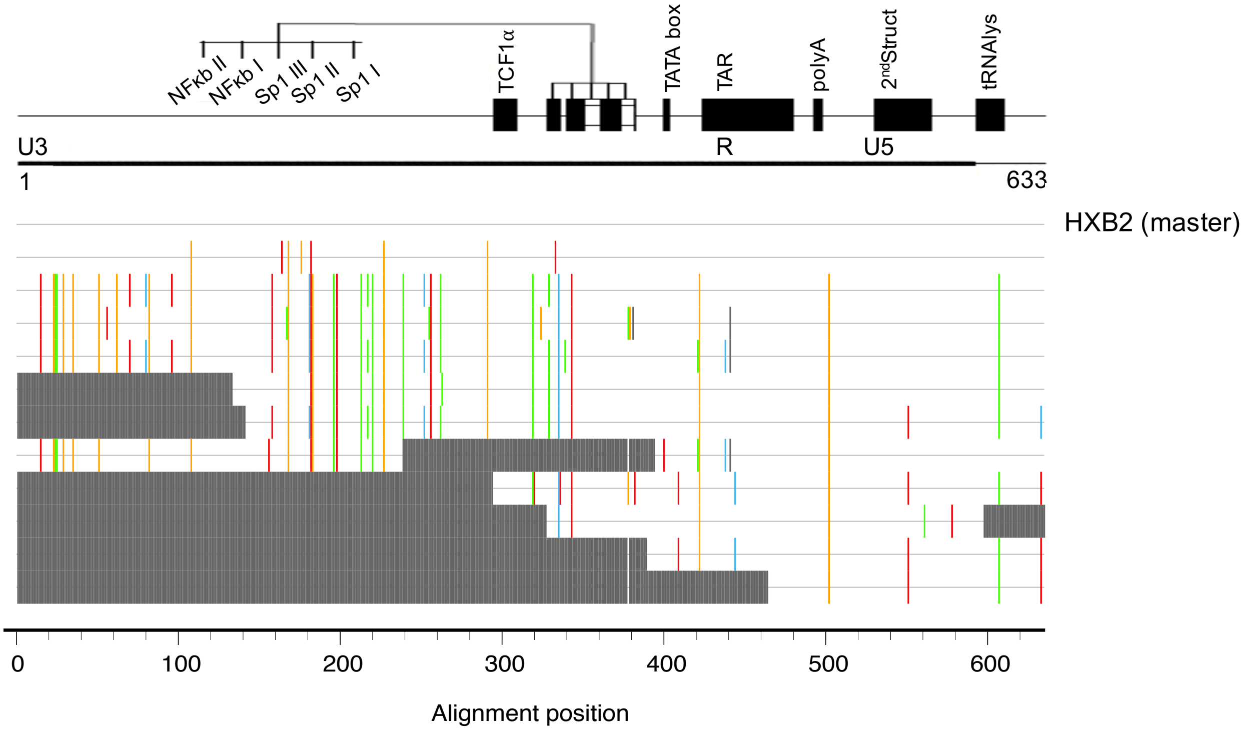


**Supplementary Figure 5.** Analysis of HIV LTR sequences in ART suppressed individual. Figure shows HIV Highlighter plots[2] of the LTR regions identified by SIP-seq in individual #1.

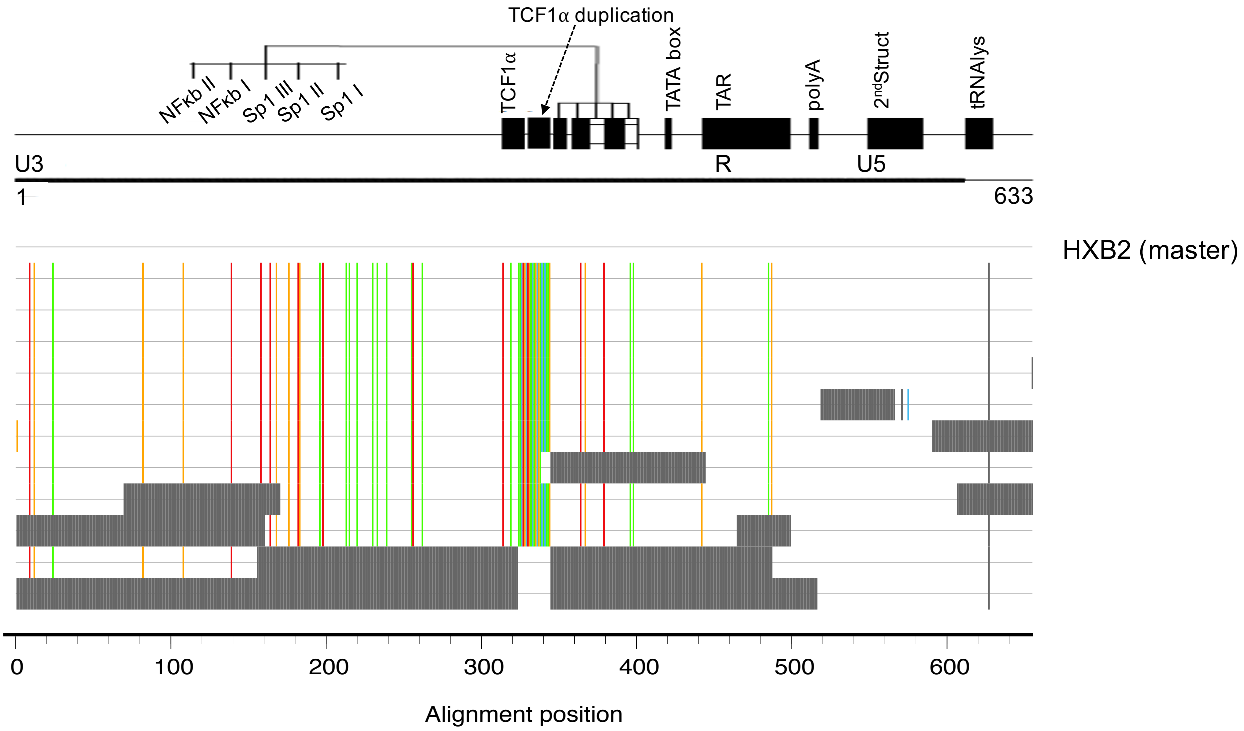


**Supplementary Figure 6.** Analysis of HIV LTR sequences in donors without ART suppression. Figure shows HIV Highlighter plots[2] of the LTR regions identified by SIP-seq in individual #3.

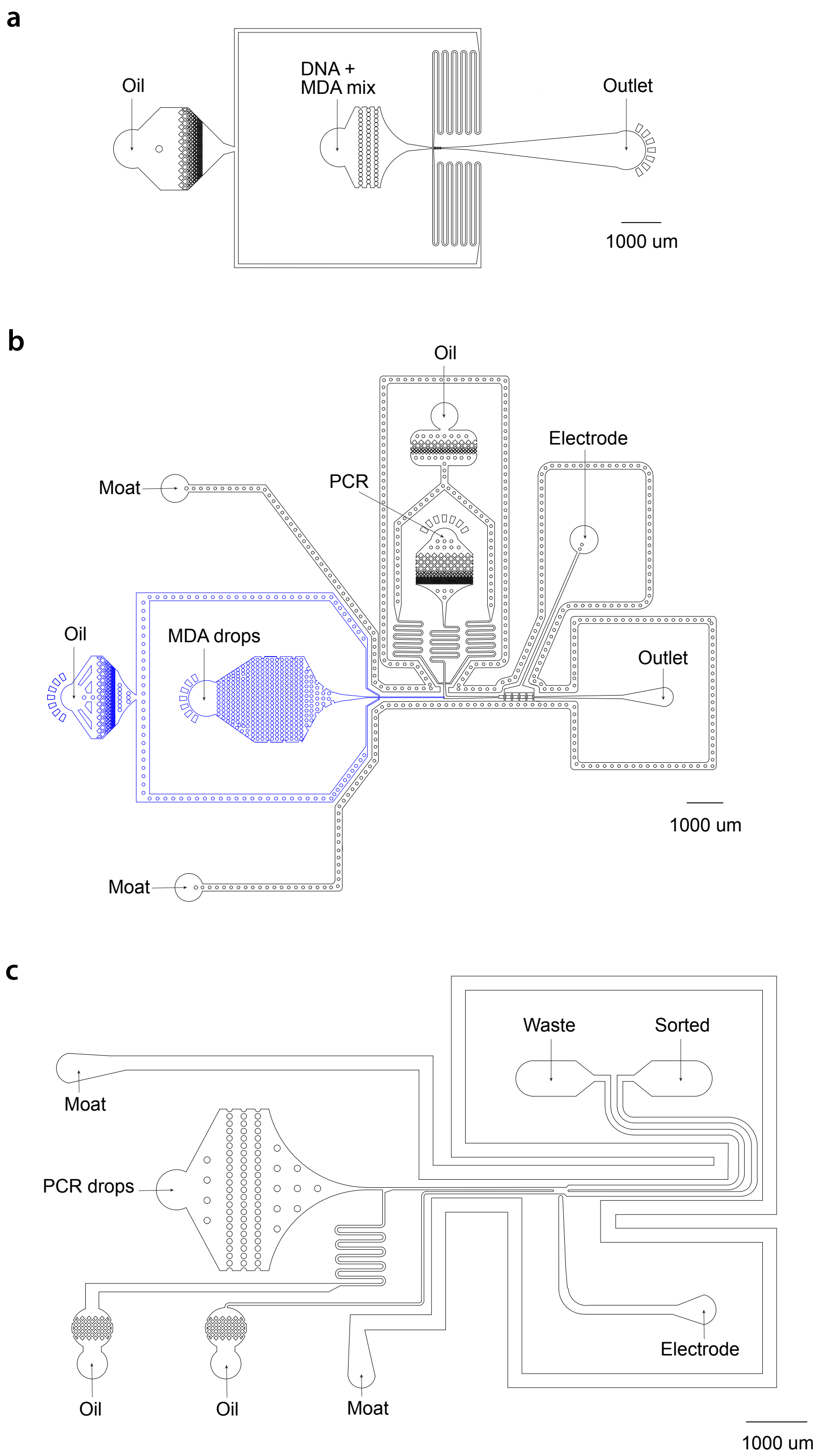


**Supplementary Figure 7.** Schematics of microfluidic devices used to: (a) encapsulate DNA and MDA reagents into isolated droplets; (b) merge MDA droplets with PCR mix droplets; and (c) sort individual PCR positive droplets.

**Supplementary Table 1: Participant Characteristics**

| Individual | Viral Load (Copies/mL) | Therapeutic Regimen |
|---|---|---|
| #1 | <20 | darunavir, ritonavir, raltegravir |
| #2 | None detected | darunavir, ritonavir, etravarine, emtricitabine, tenofovir AF |
| #3 | 102,045 | Lamiduvine, Indinavir, Stavudine |
| #4 | 833,880 | Zidovudine, Lamiduvine, Indinavir |

**Supplementary Table 2: Primers and probes for SIP-seq**

| Primer Name | HXB2 coordinates | Fluorophore, quencher | Sequence (5→3') |
|---|---|---|---|
| *pol* probe | 2586–2604 | FAM, ZEN-3'IBFQ | AAGCCAGGAATGGATGGCC |
| *pol* F | 2536-2562 | | GCACTTTAAATTTTCCCATTAGYCCTA |
| *pol* R | 2662-2634 | | CAAATTTCTACTAATGCTTTTATTTTTTC |
| *env* probe | 7815-7791 | Cy5, TAO-3'IBRQSp | CCATAGTGCTTCCTGCTGCTCCCAA |
| *env* F | 7724-7744 | | GGCAARGAGAAGAGTGGTGCA |
| *env* R | 7851-7832 | | GYCTGGCYTGTACCGTCAGC |
| LTR probe | 581-599 | VIC, 3' MGQNFQ | CTGGTAACTAGAGATCCCT |
| LTR F | 522/9607-543/9628 | | GCCTCAATAAAGCTTGCCTTGA |
| LTR R | 633/9718-608/9693 | | GCTAGAGATTTTCCACACTGACTARA |
| ψ probe | 682-705 | Cy5, TAO-3'IBRQSp | TCTCTCGACGCAGGACTCGGCTTG |
| ψ F | 633-649 | | CAGTGGCGCCCGAACAG |
| ψ R | 773-795 | | ACCCATCTCTCTCCTYCTAGCCT |
| *gag* probe1 | 1399-1416 | FAM, 3' MGQNFQ | ACCATCAATGAGGAAGCT |
| *gag* probe2 | 1430-1417 | FAM, 3' MGQNFQ | CTATCCCATTCTGC |
| *gag* F | 1372-1391 | | CAAGCAGCYATGCARATGTT |
| *gag* R | 1504-1482 | | TTCCTGCTATRTCACTTCCCCTT |
| *int* probe | 4977-4960 | VIC, 3' MGQNFQ | CTACTGCCCCTTCACCTT |
| *int* F | 4927-4947 | | CACTTTGGAAAGGACCAGCAA |
| *int* R | 5065-5044 | | TCATCACCTGCCATCTGTTTKC |
| *tat probe* | 5968-5993 | Aby, 3' QSY | CTATGGCAGGAAGAAGCGGAGACAGC |
| *tat* F | 5909-5931 | | TGTAAAAAGTGTTGCYTTCATTG |
| *tat* R | 6054-6033 | | ACTACTTACTGCTTTGRTAGAG |

**Supplementary Table 3: Primers for post SIP-seq nested PCR**

| Primer Name | Sequence (5→3’) |
|---|---|
| *ARIH2_1_outerF* | TCATGTTCCAGGCACTTTATCT |
| *ARIH2_1_outerR* | GGATCTACTGGCTCCATTTCTC |
| *ARIH2_1_innerF* | TGTTGTGAAGGAGGTGTTATCAT |
| *ARIH2_1_innerR* | CACTCTCCTCTGGCGAATAATG |
| *ARIH2_2_outerF* | GCAAGTAGACAGGATGAGGATTAG |
| *ARIH2_2_outerR* | TATGTCTGTTGGGCCTTGTC |
| *ARIH2_2_innerF* | ATGTTTCAGGGAAAGCTAGGG |
| *ARIH2_2_innerR* | TCTGGGCTAGTGGTCAGAT |
| *RABL6_1_outerF* | TGAGCTACTTGGGAGGCTAA |
| *RABL6_1_outerR* | GGGTCTACAGACAGGAGAAAGA |
| *RABL6_1_innerF* | CACTTGAGCCCAGGAGATTG |
| *RABL6_1_innerR* | AGTAGACCCTAACCTAGCAGAC |
| *RABL6_2_outerF* | CAGGGAACTAGAGCGATTT |
| *RABL6_2_outerR* | CATATCATTTACCAGCTGTCAT |
| *RABL6_2_innerF* | CGGCAGGCTGTAGACAAATA |
| *RABL6_2_innerR* | GTGGCTCACGCCTCTAATC |
